# Identification of 12 cancer types through genome deep learning

**DOI:** 10.1101/528216

**Authors:** Yingshuai Sun, Sitao Zhu, Kailong Ma, Weiqing Liu, Yao Yue, Gang Hu, Huifang Lu, Wenbin Chen

## Abstract

**Motivation:** Cancer is a major cause of death worldwide, and an early diagnosis is required for a favorable prognosis. Histological examination is the gold standard for cancer identification; however, there is a large amount of inter-observer variability in histological diagnosis. Numerous studies have shown that cancer genesis is accompanied by an accumulation of harmful mutations within patients’ genome, potentiating the identification of cancer based on genomic information. We have proposed a method, GDL (genome deep learning), to study the relationship between genomic variations and traits based on deep neural networks with multiple hidden layers and nonlinear transformations.

**Result:** We analyzed 6,083 samples from 12 cancer types obtained from the TCGA (The Cancer Genome Atlas) and 1,991 healthy samples from the 1000 Genomes project(Genomes Project, et al., 2010). We constructed 12 specific models to distinguish between certain types of cancers and healthy tissues, a specific model that can identify healthy vs diseased tissues, and a mixture model to distinguish between all 12 types of cancer based on GDL. We present the success obtained with GDL when applied to the challenging problem of cancer based on genomic variations and demonstrate state-of-the-art results (97%, 70.08% and 94.70%) for cancer identification. The mixture model achieved a comparable performance. With the development of new molecular and sequencing technologies, we can now collect circulating tumor DNA (ctDNA) from blood and monitor the cancer risk in real time, and using our model, we can also target cancerous tissue that may develop in the future. We developed a new and efficient method for the identification of cancer based on genomic information that offers a new direction for disease diagnosis while providing a new method to predict traits based on that information.

## 1 Introduction

Cancer is the most common risk that threatens human health worldwide. There are more than 100 types of cancers, including cancers of the breast, skin, lung, colon, prostate and ovaries. In the United States, 1,735,350 new cancer cases and 609,640 cancer deaths will be reported in 2018(Siegel, et al., 2018). It is known that cancer is mainly caused by harmful mutations in proto-oncogenes, tumor suppressor genes and cell cycle regulator genes. For example, previous studies indicated that *p53* activates DNA repair proteins and inhibits the occurrence of various types of cancer(Olivier, et al., 2010). In breast cancer, high penetrance mutations in *BRCA1* and *BRCA2* cause a loss of tumor suppressive function which correlates with an increased risk of breast cancer(Petrucelli, et al., 2010). In addition, *C21orf58* and *ZNF526* also have functional roles in the control of breast cancer cell growth(Zhang, et al., 2018). Although stomach cancer is caused by infection with the bacteria *H. pylori*, the genetic basis of this cancer remains largely unknown. There are published reports that stomach cancer may be caused by the accumulation *PBLB2* and *ATM* mutations(Hannes, 2015). BLCA (Bladder Urothelial Carcinoma) is a major cancer of the urinary system. TCGA researchers have identified many mutated genes that are involved in the cell cycle, DNA repair and chromatin modifications in BLCA. *BLCA-4*(Myers-Irvin, et al., 2005), a nuclear matrix protein, plays a major role in bladder cancer carcinogenesis. Although many genes that have been found have major roles in the occurrence and spread of cancer, the pathogenic mechanisms of gene mutations and interactions between genes are largely unknown. In this work we studied twelve cancer types including BLCA, BRCA (breast adenocarcinoma), COAD (colon adenocarcinoma), GBM (glioblastoma multiforme), KIRC (kidney renal clear cell carcinoma), LGG (low grade glioma), LUSC (lung squamous cell carcinoma), OV (ovarian carcinoma), PRAD (prostate adenocarcinoma), SKCM (skin cutaneous melanoma), THCA (thyroid carcinoma) and UCEC (uterine corpus endometrial carcinoma) that are affected by genomic variations that are distributed widely across the genome (Figure 1). With the development of DNA sequencing and bioinformatics analysis methods, we have been able to identify additional genomic mutations and have accumulated a large amount of data. Methods for identifying correlations between mass genomic variations and cancer are urgently required.

**Figure 1.**
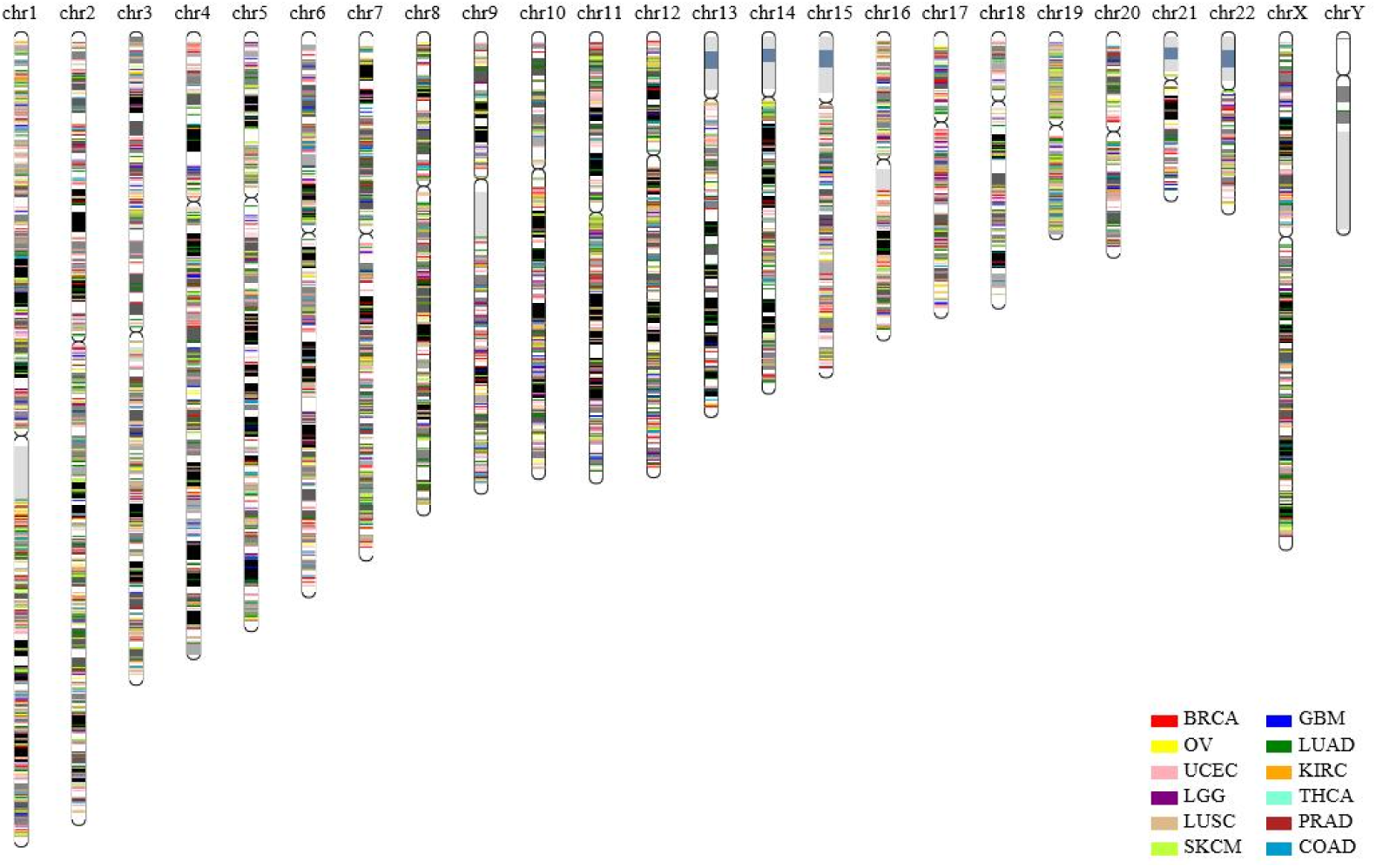
The distribution of 12 cancer point mutations in the entire genome. The chromosome block was produced a visible karyotype by staining condensed chromosomes (Giemsa banding). Adenine and thymine (AT) rich and relatively gene-poor area was stained more darkly. Guanine and cytosine (GC) rich and more transcriptionally active area was stained lightly. The legend certain color represent the cancer in the lower right corner.

Deep learning algorithms, such as Alpha Go(Silver,et al.,2016) and object recognition(Russakovsky, et al., 2015), exceed human performance in visual tasks and are flexible and powerful analytical techniques for dealing with complex problems. Deep learning is a high-level abstraction algorithm for solving classification and regression problems. Through deep learning and pattern mining of data, it identifies complex structures in massive data sets and has great potential for applications in genetics and genomics(Libbrecht and Noble, 2015). As a novel technique, a number of cases were shown to provide better performance in biological applications. Deep learning algorithms can be used to learn how to recognize the locations of splice site promoters and enhancers(Cuperus, et al., 2017). Deep learning algorithms also have many applications in the prediction of protein secondary structure and function(Li, et al., 2017). More accurate identification of phenotypes would improve study efficiency through a convolutional neural network(Pound, et al., 2017), which is one image recognition algorithm of deep learning algorithms. Researchers also found that the skin cancer identification rate using deep neural networks was more accurate than that determined by dermatologists(Esteva, et al., 2017). Deep learning algorithms use multiple layers of nonlinear processing units for feature extraction and transformation to find deep relationships between complex variations under supervised or unsupervised procedures(LeCun, et al., 2015).

Biological traits are the result of interactions between gene sequences and gene interactions under certain environmental conditions. The deep learning model is suitable for studying the relationship between these factors and the phenotype. We constructed a model for the identification of cancer based on genomic variations that we call “genomic deep learning” (GDL). GDL studies the relationship between genomic variations and traits through deep learning of genomes. At present, cancer research is mainly based on genome-wide association analysis (GWAS). GWAS is meant to identify associations between single nucleotide variations and cancer(Hirschhorn and Daly, 2005). GWAS is based on linkage analysis to find the diseased genes and requires more intimate segregate sites(Hirschhorn and Daly, 2005). However, deep learning models can take entire genome variations into account without the influence of segregate sites. Deep learning is an algorithm that simulates biological neural networks. Neural network algorithms are similar to biological trait regulation networks. It is possible and feasible to build a deep neural network (DNN) model for the identification of cancer via massive variants.

In this work we constructed 12 specific, a total-specific and mixture cancer identification models using a deep neural network (DNN) within a TensorFlow(Rampasek and Goldenberg, 2016) framework. We used an exponential decay method to optimize the learning rate, L2 regularization(Mocanu, et al., 2018) to minimize overfitting, and a sliding average model to increase the robustness of the model. For each specific model meant to identify a certain type of cancer, the detection accuracy, sensitivity and specificity can be greater than 97%, 98% and 97%, respectively. The mixture model, which is able to distinguish all 12 types of cancers, exhibited comparable performance. The total-specific and mixture models also demonstrated comparable performance. The development of molecular technologies enables us to collect circulating tumor DNA(Abbosh, et al., 2018) (ctDNA) from blood. Using our model, cancerous tissue can be identified more conveniently and timely, thus providing an opportunity for earlier treatment. This approach to genome deep learning offers a new direction for disease diagnosis while providing a new method to predict traits based on genomic information.

## 2 Genome Deep Learning Methodology

Cancer is caused by the accumulation of harmful mutations(Wishart, 2015). Mutations occur all the time, especially during cell genome duplication, but most of the mutations are not on key genes. If the harmful mutations occur in the oncogene or tumor suppressor genes, the normal cells will become cancer cells. Changes in multiple genes are required to transform a normal cell into a cancer cell. To determine the relationship between mutations and cancers, we designed a deep learning method that we call genomic deep learning (GDL). GDL is a classification method for cancer identification. The architecture of our model contains feature selection, feature quantization, data filters and deep neural networks involving multiple hidden layers between input and output layers (Figure 2). Multiple hidden layers are used to simulate the process of genome expression.

**Figure 2.**
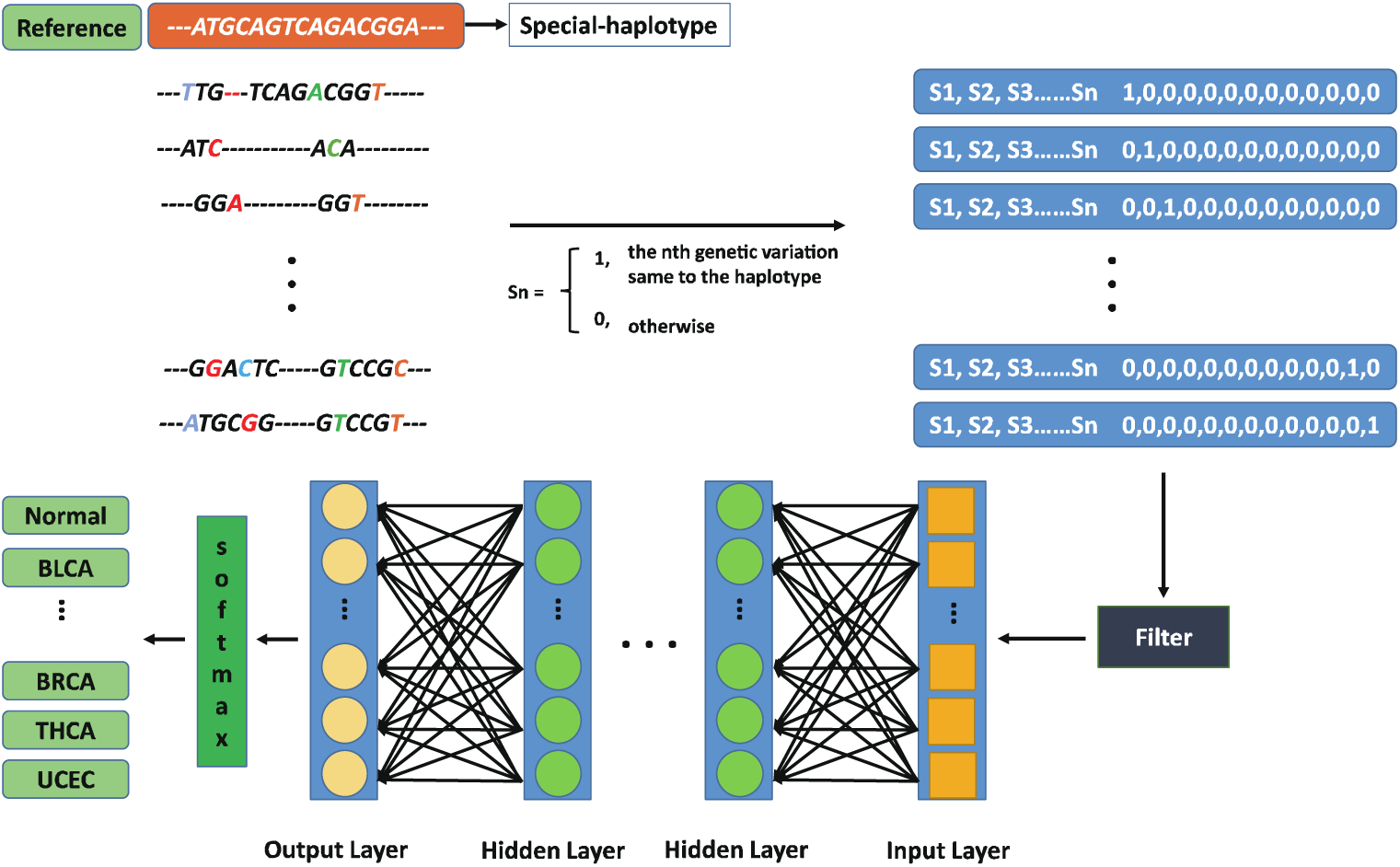
The architecture of genomic deep learning (GDL). Firstly, the special haplotype was obtained in orange box. Secondly, control and treatment were transformed according to the variation files. Finally, the filtered data was put into the GDL model, training step by step.

GDL consists of data processing and model training. Data processing consists of three steps. First, the sequencing data are compared with a reference to obtain a point mutation file, and then the point mutation file is converted into a format of the model input. The third step is to filter the data after conversion formatting. Model training is DNN modeling and includes an input layer, multiple hidden layers, an output layer and a softmax layer. After training, a classification model is finally obtained.

### 2.1 Data processing

#### 2.1.1 Model feature selection and quantification

The types of genetic mutations that cause cancer include point mutations, insertion or deletion (indel) mutations, structural variations (SV) and copy number variations (CNV). SVs and CNVs are difficult to quantify accurately; however, point mutations and indel mutations can be accurately quantified, and they have the highest prevalence and accuracy. Therefore, we chose point mutations and indel mutations to construct our model.

It is obviously impractical to select all of the point mutations as dimensions for the model because mass dimensions will increase the computation cost. To reduce the learning pressure brought about by highly redundant dimensions and to reduce the learning difficulty without affecting the accuracy of the model, we selected point mutations closely related to cancer from the TCGA as the dimension for the model. In specific models, we ranked the point mutations of each cancer according to the number of occurrences in this cancer group from high to low. We choose different former points as the dimensions (1k, 2k, 3k, 4k, 5k, 6k, 7k, 8k, 9k and 10K) for modeling. The results demonstrate that with an increase of the dimension, the accuracy will continue to improve (Supplementary Fig. 2). Finally, we chose 10,000 point mutations as the dimension for the specific models. The accumulation of harmful mutations is the root cause of cancer. In the development of cancer, the accumulation of mutations can be divided into two parts: the first part is the accumulation of mutations that occur that lead to the cancer, and the second part is the accumulation of mutations that occur after the cancer develops, which is the cause of tumor heterogeneity. Our goal was to determine the rules that gene mutations follow in converting healthy tissues to cancer, which is reflected in the effect of the mutations on the pathways involved. The difference in genetic mutations between patients with the same type of cancer is large because the effect on different pathways is similar. The second part of the mutation occurs on the basis of the first part of the variation. To determine these rules faster using limited computing resources, we choose the position where the number of occurrences of the variation is greater than or equal to 2 as the dimension in the total-specific model and the mixture model.

Point mutations are used as model input for GDL. The Edico Genome Pipeline reduces the time required for analyzing an entire genome at 30x coverage from ∼10 hours (BWA and GATK software)(Li and Durbin, 2009; McKenna, et al., 2010) and was chosen to use point mutations as a baseline for healthy tissues. The reference genome was GRCh38, which was downloaded from the National Cancer Institute website (https://gdc.cancer.gov/about-data/data-harmonization-and-generation/gdc-reference-files). Mutect2, a method that applies a Bayesian classifier to detect somatic mutations with very low allele fractions, requires only a few supporting reads, followed by carefully tuned filters to ensure that high specificity was used for calling cancer point mutations(Cibulskis, et al., 2013).

In any application of deep learning methods, the researcher must decide which data format to provide as input to the algorithm. How to convert VCF data into GDL model input format becomes a significant challenge. To overcome this input format challenge, the HapMap(International HapMap, 2005) project provided us with an approach. High risk sites were collected form the TCGA(Ding, et al., 2018) and we then sorted the collected sites by the frequency of occurrence in cancer patients named as a “special-haplotype”. Furthermore, variant sites from healthy people and cancer patients were assigned a score (“0” indicates different from special-haplotype and “1” indicates the same as special-haplotype) compared to the special-haplotype. Finally, our input file became an array, for example: 1,1,0,0….1,0. As described above, our variant format was transformed into a different classifier, healthy or cancer. Each type of situation has its own classification label which is expressed by One-Hot Encoding. For example, the class labels “1,0,0,0,0,0,0,0,0,0,0,0” and “0,1,0,0,0,0,0,0,0,0,0,0” represent BLCA and BRCA, respectively. Other cancer types were treated as described above. Finally, the VCF files were transformed into two parts separated by a space. In part one, Sn had only two choices. A “1” indicates that the special individual genomic variation was the same as the special-haplotype, and a “0” indicates that they were different. Sn represents each genomic variation in an n index. In part two, the class label indicates whether the individual was healthy or not (Figure 2).

#### 2.1.2 Dataset and data filter

To collect point mutations for the DNN model we used VCF files containing genomic variations between paired healthy tissues from the IGSR (The International Genome Sample Resource) and tumor tissues from the TCGA entire exon sequencing (WES) database. Meanwhile these datasets can also be availabled in CNGBdb. The involved tumor tissues were from twelve cancer types including BLCA, BRCA, COAD, GBM, KIRC, LGG, LUSC, OV, PRAD, SKCM, THCA and UCEC. Each type of tumor tissue was comprised of 425, 1080, 493, 498, 376, 530, 561, 610, 503, 472, 504 and 561 samples for a total of 6083 datasets. We also downloaded genome sequencing data from 1,991 healthy individuals from the 1000 Genomes Project database (http://www.internationalgenome.org/).

Due to the limitations of the amount of data, the feature dimensions for each cancer cannot be fully selected, and therefore existing dimensions may not be able to completely distinguish different cancers. To prevent this situation from negatively affecting the modeling process, we filtered the data. First, if the cancer patient had no variants in the special-haplotype reference, that patient would be removed. Second, duplicate healthy individuals should be unique.

### 2.2 Training

#### 2.2.1 DNN model function

The DNN model was composed of several computational layers. Each layer takes an input and produces an output, often computed as a non-linear function of weighted linear combinations of the input layer and adjusts each weight and threshold by accumulated error back propagation. In the forward propagation process, the output from each neuron is a nonlinear calculation of the weighted sum of the previous layer pointing to that neuron(Schmidhuber, 2015). The formula used is

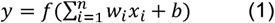

where y represents the output of activation, n represents the number of hidden units in the layer, *w* and *x*_*i*_ are the input of the activation, and b represents the bias terms.

Activation functions play an important role in deep learning because the combination of arbitrary linear models is still a linear model. To solve more complex problems, we used the activation function to achieve de-linearization. The commonly used activation functions are ReLU(Nair and Hinton, 2010), Sigmoid and Tanh. The calculation using Sigmoid is relatively complex and requires a very long running time, and the gradient is easy to lose during the process of back propagation. Tanh also requires a large amount of calculation time. Although ReLU is relatively fragile, it requires a relatively small amount of computation, and it has faster convergence speed. The other advantage was that ReLU causes sparsity of the network and reduces interdependence of parameters that overcome the occurrence of overfitting problems. Formula 2 is the formula for the ReLU function:

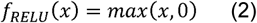

After completing the current propagation, we use the loss function to represent the difference between the predicted and target values to evaluate the model’s effectiveness. The process of training the model is the process of decreasing the loss function. After the hidden layer of the model, the output of the hidden layer becomes a probability distribution through the softmax layer. We then use the cross entropy as a loss function to calculate the distance between the predicted probability distribution and the true probability distribution.

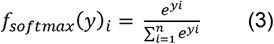

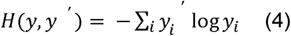

#### 2.2.2 Model optimization

To obtain a better model, we further optimized it based on back propagation(Rumelhart, et al., 1986) and gradient descent. In the model training process, the learning rate controls the speed of the model update. If the learning rate is too high, the parameters will move back and forth on both sides of an acceptable value. On the contrary, if the learning rate is too small, convergence can be guaranteed, but the speed of optimization will be greatly reduced. Therefore, we used a more flexible learning rate setting method, i.e., exponential decay. With this method, a relatively large learning rate can be used to obtain a better result more quickly, and the learning rate is then gradually reduced with subsequent iterations, making the model more stable in the later period of training. Formula 5 is the formula for the exponential decay of the learning rate, where R represents the decayed learning rate, r represents the basic learning rate, d represents the decay rate, g represents the global step, and s represents decay step. Due to sequencing errors and the limitations of obtaining point mutation algorithms, false positive and false negative data are unavoidable in our data. If the model can remember the noise in each training data well, it will forget to learn the general trend in the training data. We use L2 regularization as an index of model complexity, and then add it to the loss function to reduce the model complexity and avoid overfitting problems. Formula 6 is the formula for the L2 regularization, where w_i_ represents weights. To improve the robustness of the model in the test data, we use the sliding average model which can reduce periodic interference and effectively remove the random fluctuations in the prediction. This approach maintains a shadow variable for each variable, and each time the variable is updated, the independent variable is also updated. Formula 7 is the formula for the shadow variable, where S represents the shadow variable, d represents decay and V represents a variable.

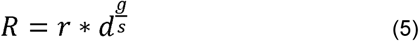

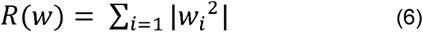

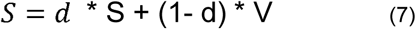

The DNN model was implemented in TensorFlow(Rampasek and Goldenberg, 2016) and the Google open source software library using data flow graphs and was trained on a Mac OS. In TensorFlow, networks are constructed and executed in a TensorFlow graph. Twelve cancer types, abbreviated as BRCA, OV, UCEC, LGG, LUSC, SKCM, GBM, LUAD, KIRC, THCA, PRAD and COAD were chosen to construct the DNN model.

#### 2.2.3 Model explanation

The DNN module in the GDL method is the key to computer learning of gene expression. In its hidden layers, we gradually abstract the advanced features and then summarize the three stages according to the biological processes involved. In the first stage, a gene mutation is abstracted according to the directional variation of the fixed position in the input layer. It is beneficial that according to the combination of different weights in the input layer, the first stage can simulate the different effects of different combinations of variations on genes and can even simulate alternative splicing. Alternative splicing occurs as a normal phenomenon in eukaryotes, where it greatly increases the biodiversity of proteins that can be encoded by the genome(Black, 2003). Abnormal splicing variants are also recognized to be highly associated with cancer(Fackenthal and Godley, 2008; He, et al., 2009; Skotheim and Nees, 2007), and splicing factor genes are frequently mutated in different types of cancers(Sveen, et al., 2016). In the second stage, the first stage of gene mutation is further abstracted as a variation of the pathway. Genetic alterations in signaling pathways that control cell-cycle progression, apoptosis and cell growth are common hallmarks of cancer. In the third stage, the variation information of all the pathways in the second stage is further abstracted as a manifestation of cancer. Those features described above would help us understand the model for the biological process. Nevertheless, deep neural networks can abstract more features from higher dimensions. In addition to those visible features, the DNN model can abstract invisible features.

### 2.3 Evaluation

Model evaluation produces an intuitive understanding of model reliability. To identify each cancer type, since it is a binary classification, we use accuracy, sensitivity and specificity to evaluate the classifiers’ performance. Since the total DNN model is a multi-class classification problem, we use accuracy, sensitivity and specificity to evaluate the total classifiers’ performance. Sensitivity, specificity and accuracy of the classification were calculated using results from all validation subsets. After the softmax function, if the probability score for a cancer was higher than the threshold value, the predictive diagnosis was a special cancer type.

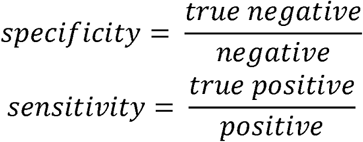

## 3 Results

### 3.1 Cancer types and samples statistics

Genomic variation files for healthy people (1991) and cancer patients (6083) were obtained from the 1000 Genome Project Website and the TCGA online database. As shown in Table 1, the sample number of each tumor type ranges from 339 (KIRC) to 1,044 (BRCA). From 6,083 TCGA samples with available information, 71.61% (n = 4,356) were White, 9.73% (n = 592) were American, 3.98% were Asian and 14.68% have no race information. Patients (n = 6,083) were diagnosed between 55 and 74 years of age. The sex distribution had no serious effect, except for prostate adenocarcinomas (PRAD), which were all male, and ovarian carcinomas (OV) which were all female. Cancer staging plays an important role in determining treatment options. According to the cancer stage standards, all cancers were divided into four stages and one unidentified stage. From Table 1, we can see that GBM, LGG, OV, PRAD and UCEC have no clear stage. Sequencing reads of 1991 samples from the 1000 Genome project were analyzed using the Edico genome pipeline. We obtained 25 Tb of next-generation sequencing (NGS) data from the mainstream sequencing platforms including Illumine and Solid. The sequencing depth for healthy people ranged from 4 to 10 (Supplementary Table1).

**Table 1.**
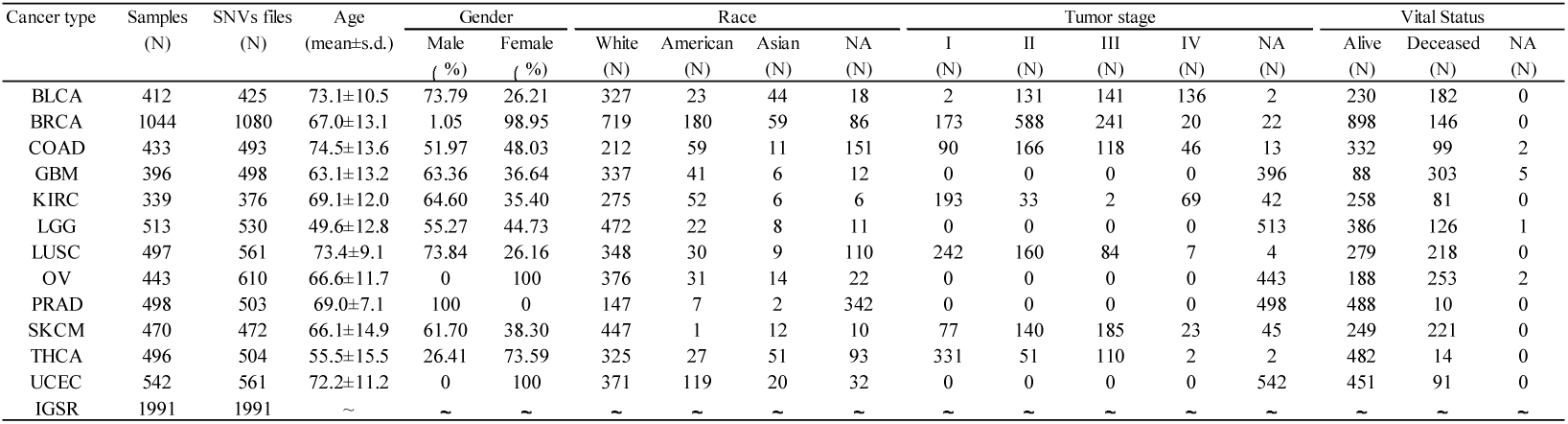
Summary information of datasets from the TCGA and the 1000 Genome Project that were used in this study.

### 3.2 Accuracy of cancer identification

After an extended period of data preparation and model training, an acceptable classification result was obtained. All specific models showed accuracy ranges from 97.83% to 100% (mean 98.89%, Figure 3), with sensitivity ranges from 97% to 100% and specificity ranges from 98% to 100% (Table 2). The accuracy, sensitivity and specificity of the total-specific model were 94.7%, 97.3% and 85.54%, respectively. We used ROC and AUC to evaluate the direct performance of the specific models (Figure 4a). Each model exhibited a high AUC and was completely correct in four models, i.e., BLCA, KIRC, OV and THCA. Such high quality classification models demonstrate that significant differences exist between patients and healthy people (Supplementary Figures 4-6).

**Table 2.** Summary of the GDL model classification performances.

| Cancer type | Raw data |  | Filter |  |  |  |  |  | Accuracy<br>(%) | Sensitivity<br>(%) | Specificity<br>(%) |
| --- | --- | --- | --- | --- | --- | --- | --- | --- | --- | --- | --- |
|  |  |  | ALL |  | Train Data |  | Test Data |  |  |  |  |
|  | Cancer | Health | Cancer | Health | Cancer | Health | Cancer | Health |  |  |  |
| BLCA | 425 | 1991 | 417 | 216 | 341 | 165 | 76 | 51 | 98.43 | 98.68 | 98.04 |
| BRCA | 1080 | 1991 | 1073 | 586 | 856 | 471 | 217 | 115 | 98.19 | 97.70 | 99.13 |
| COAD | 493 | 1991 | 482 | 842 | 385 | 675 | 97 | 167 | 99.24 | 98.97 | 99.40 |
| GBM | 498 | 1991 | 478 | 435 | 385 | 345 | 93 | 90 | 97.81 | 96.77 | 98.89 |
| KIRC | 376 | 1991 | 372 | 189 | 288 | 161 | 84 | 28 | 100.00 | 100.00 | 100.00 |
| LGG | 530 | 1991 | 518 | 491 | 410 | 397 | 108 | 94 | 99.01 | 99.07 | 98.94 |
| LUSC | 561 | 1991 | 545 | 166 | 436 | 133 | 109 | 33 | 100.00 | 100.00 | 100.00 |
| OV | 610 | 1991 | 600 | 176 | 481 | 140 | 119 | 36 | 100.00 | 100.00 | 100.00 |
| PRAD | 503 | 1991 | 494 | 497 | 399 | 394 | 95 | 103 | 97.47 | 95.79 | 99.03 |
| SKCM | 472 | 1991 | 434 | 409 | 358 | 316 | 76 | 93 | 98.22 | 97.37 | 98.92 |
| THCA | 504 | 1991 | 503 | 241 | 405 | 190 | 98 | 51 | 97.99 | 97.96 | 98.04 |
| UCEC | 561 | 1991 | 549 | 1217 | 446 | 967 | 103 | 250 | 98.02 | 98.06 | 98.00 |
| TOTAL | 6613 | 1991 | 5733 | 1629 | 4585 | 1304 | 1148 | 325 | 94.70 | 97.30 | 85.54 |

**Figure 3.**
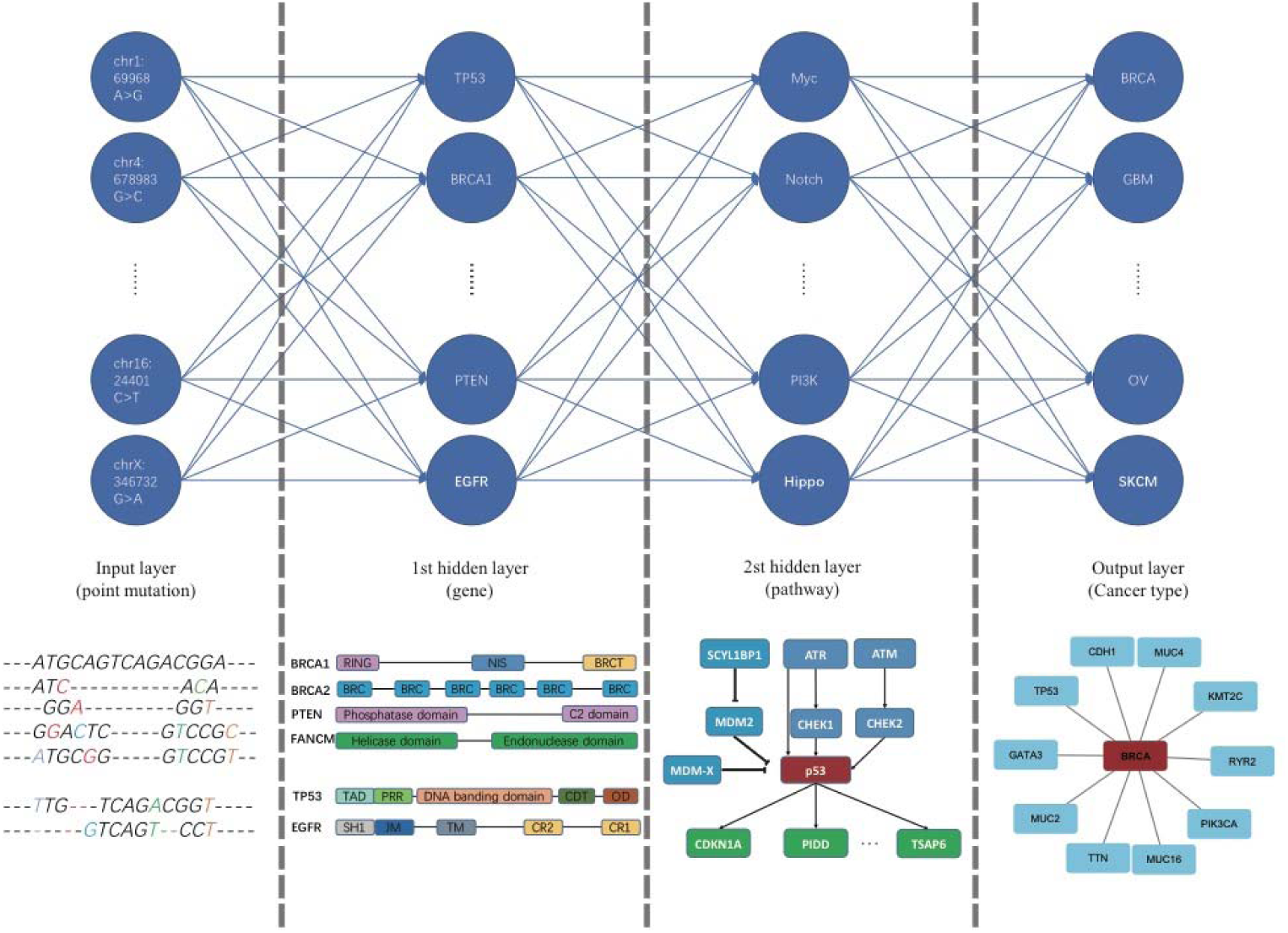
The output of each layer of a DNN in the GDL method. Abstract features were obtained from different neuron layers. The input layer, 1st hidden layer, 2nd hidden layer and output layer represent point mutation features, gene element features, the pathway feature and cancer feature, respectively.

**Figure 4.**
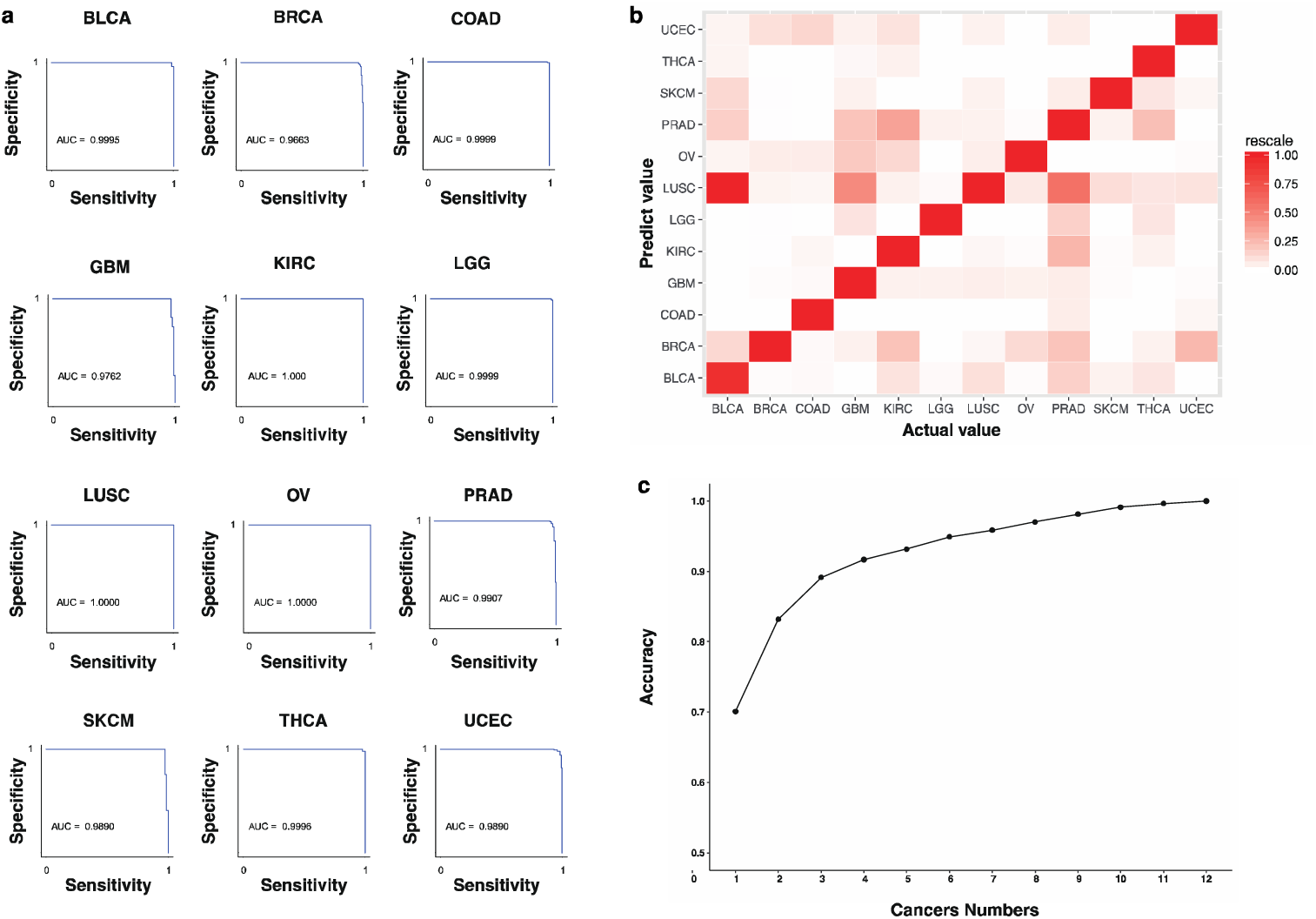
Cancer identification performance of 12 specific models and the mixture model. (a) The classification performance of 12 specific models. Using different thresholds, the sensitivity is the abscissa and the specificity is the ordinate, resulting in 12 ROC curves. The 12 ROC curves produce perfect classification results, and the area under the ROC curve (AUC) is greater than 96%. (b) Confusion matrix of the mixture mode. The abscissa indicates the label, and the ordinate indicates the predicted cancer type. LUSC is more obvious in the predictions, especially in the BLCA predictions, suggesting that many cancers are easily confused with LUSC. Cancers that are easily confused in model predictions may be similar in their genetic variations. (c) The accuracy of top-N at different forecasted quantities. The abscissa indicates different prediction numbers, and the ordinate indicates accuracy. The accuracy of the prediction result is 70.08%, and the accuracy of two prediction results is 83.20%, which provides support for the practical application of the model. The abscissa indicates the label, and the ordinate indicates the predicted cancer type.

To evaluate the model in a different aspect, we validated the DNN model using a four-cancer classification of cancer types according to the criteria staging system (tumor, node and metastasis, TNM) described in the AJCC Cancer Staging Manual^34^. The stage of the cancer is a key factor for determining the prognosis and will assist the doctor in determining the appropriate treatment. According to the criterion described in AJCC Cancer Staging Manual^34^, cancers can be divided into five levels based on the degree of tumor differentiation. In the first level (I level), the tumor has low pathogenicity and only occurs in specific areas such that the tumor has a better chance of being cured. In the fourth level (IV level), the tumor has a high degree of malignancy and has spread to other organs such that the tumor has a low probability of being cured. The last level does not meet the cancer staging standards described in the AJCC and is labeled as “None” because it is difficult to distinguish using the TCGA. For training models that use the cancer stage database, the mean accuracy for the DNN model is 97%, and the mean sensitivity and specificity is 98% and 97%, respectively (Table 3). Finally, for the mixture model, we used the data from each cancer type class to validate the DNN model.

To avoid the limitation of the specific model, we constructed a mixture model to distinguish all 12 types of cancer. The model is able to predict cancer with an accuracy of 70.08%, which is lower than that for the specific model. The accuracy of the mixture model is lower than with the specific cancer model, which is acceptable because it is a different cancer, and there is a great deal of similarity at the molecular level, causing the classification to be inaccurate(Ding, et al., 2018). Within the 12 cancers, the statistics suggested that the difference in the frequency of base mutations between different cancers is not very large. It was further demonstrated that although cancer tissues vary in form, there are large common genomic variations at the molecular level that lead to lower accuracy in the mixture model than in the specific model. Furthermore, the selection of reference sites is based on the frequency of sites in the cancer population. A high frequency of reference sites could promote better accuracy for multiply classifications in the mixture model.

To confirm a correlation between the number of common dimensions between different cancers and the judgment error in the mixed model we performed statistical analyses (Figure 4.b, Supplementary Figures 3 and 5). In Figure 5, we can see that the common dimension between UCEC and COAD is much larger than that between other cancer types. The common dimension of other groups (UCEC and BRCA, BRCA and COAD) is also higher than that between other cancer types, but much smaller than the common dimension between UCEC and COAD. However, as can be seen in Figures 2 and 3, the ratio of false judgments between UCEC and COAD is lower than that between UCEC and BRCA, which indicates that the common dimensions have no correlation with model false judgments. The ratio of false judgments between BRCA and COAD is much lower than that between the two cancers in the common dimensions.

**Figure 5.**
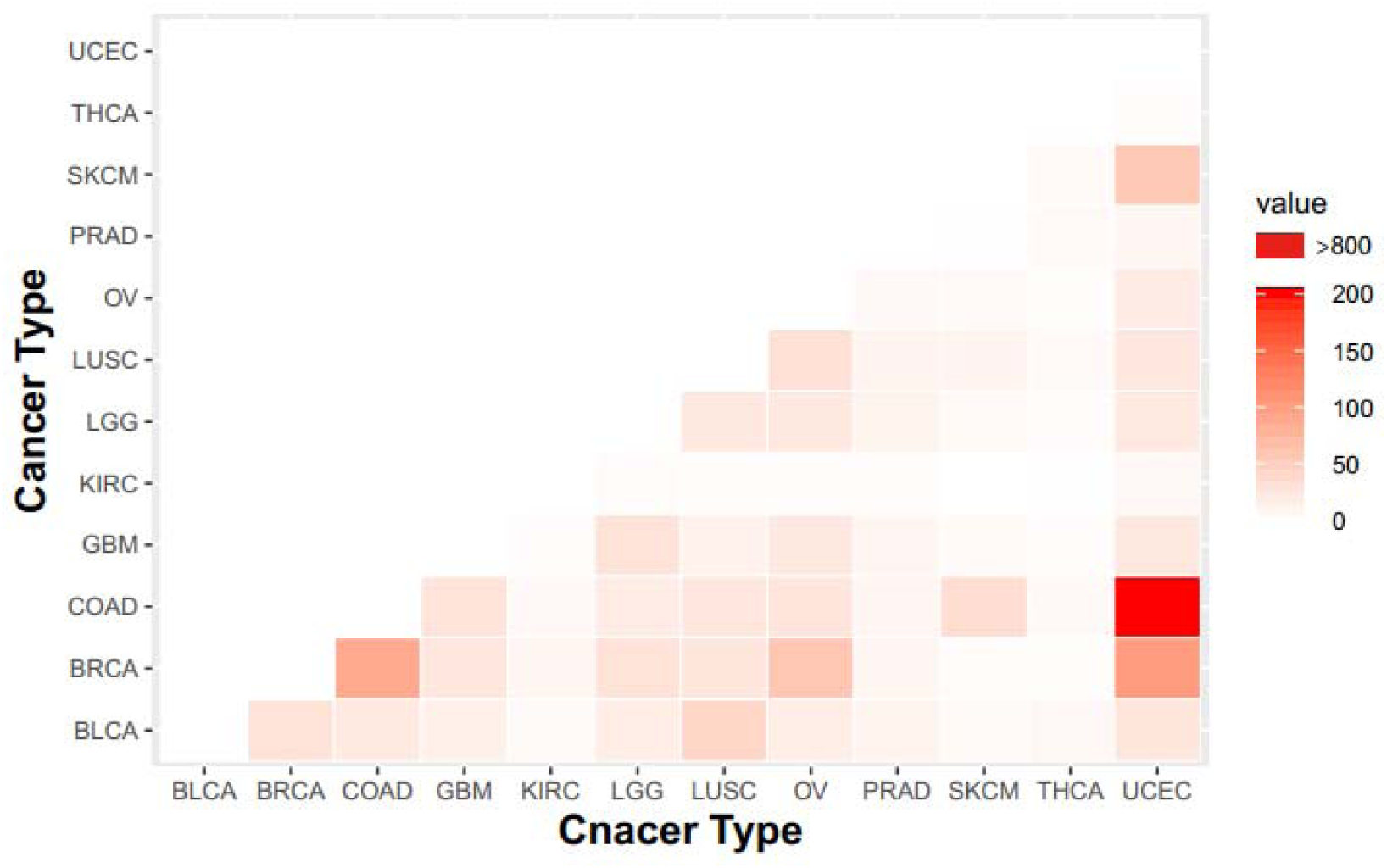
Mixed matrices of the same dimensions for different cancers. UCEC and COAD share the largest number of variant sites, followed by UCEC and BRCA. BRCA and COAD are relatively more common than other types of cancer.

## Supporting information

supplement table in main text

supplement figure in main text

## Acknowledgements

This work was supported by the bioinformatics department of BGI research, China National GeneBank, BGI-Shenzhen. We also acknowledge The Cancer Genome Atlas and 1000 Genome Project as the source of primary data. We thank Kun Ma and Wenbin Chen for helpful discussions and comments.

## Supplementary information

Supplementary data are available at Bioinformatics online.

