## supplement figure in main text for "Identification of 12 cancer types through genome deep learning"

**Supplementary Information**

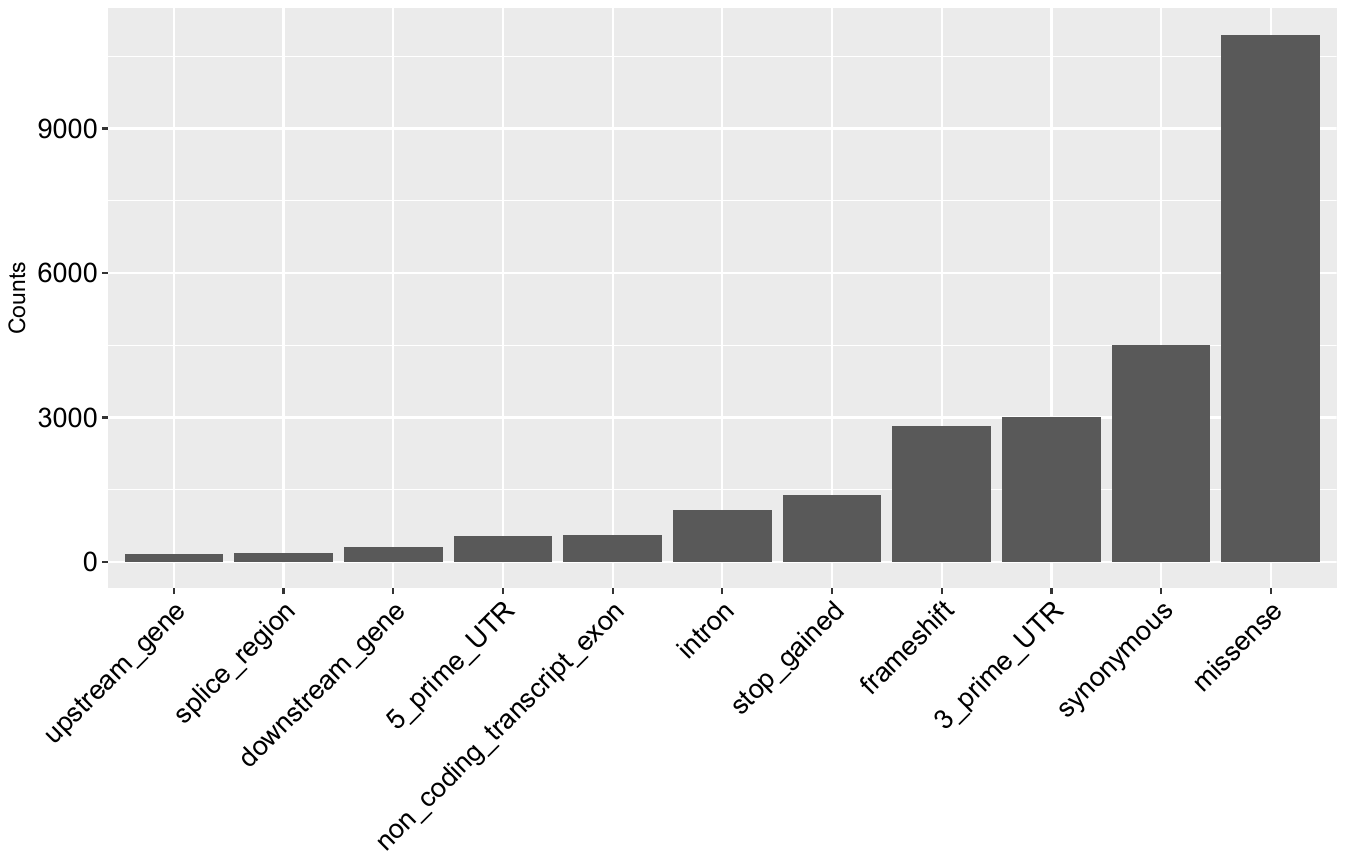

**Supplementary Figure 1 | Counts of point mutation in gene component including up stream, down stream, splice, UTR, exon and intron regions. The missense mutation are twice of synonymous mutation in counts.**

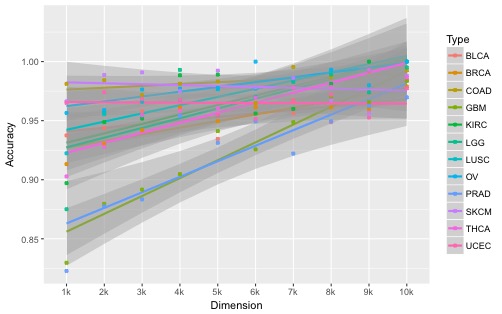

**Supplementary Figure 2 | Modeling accuracy of 12 kinds of cancer in different dimensions.**

The trends in modeling accuracy of 12 cancers in different dimensions. The accuracy of the model increases as the dimension increases. Due to the influence of training data selection and modeling parameters, the accuracy rate changes will produce some fluctuations, but from the whole trend, the accuracy rate increases with the increase of the dimension.

**Supplementary Table 1 | The performance of cancer stage dataset for GDL model evaluation.**

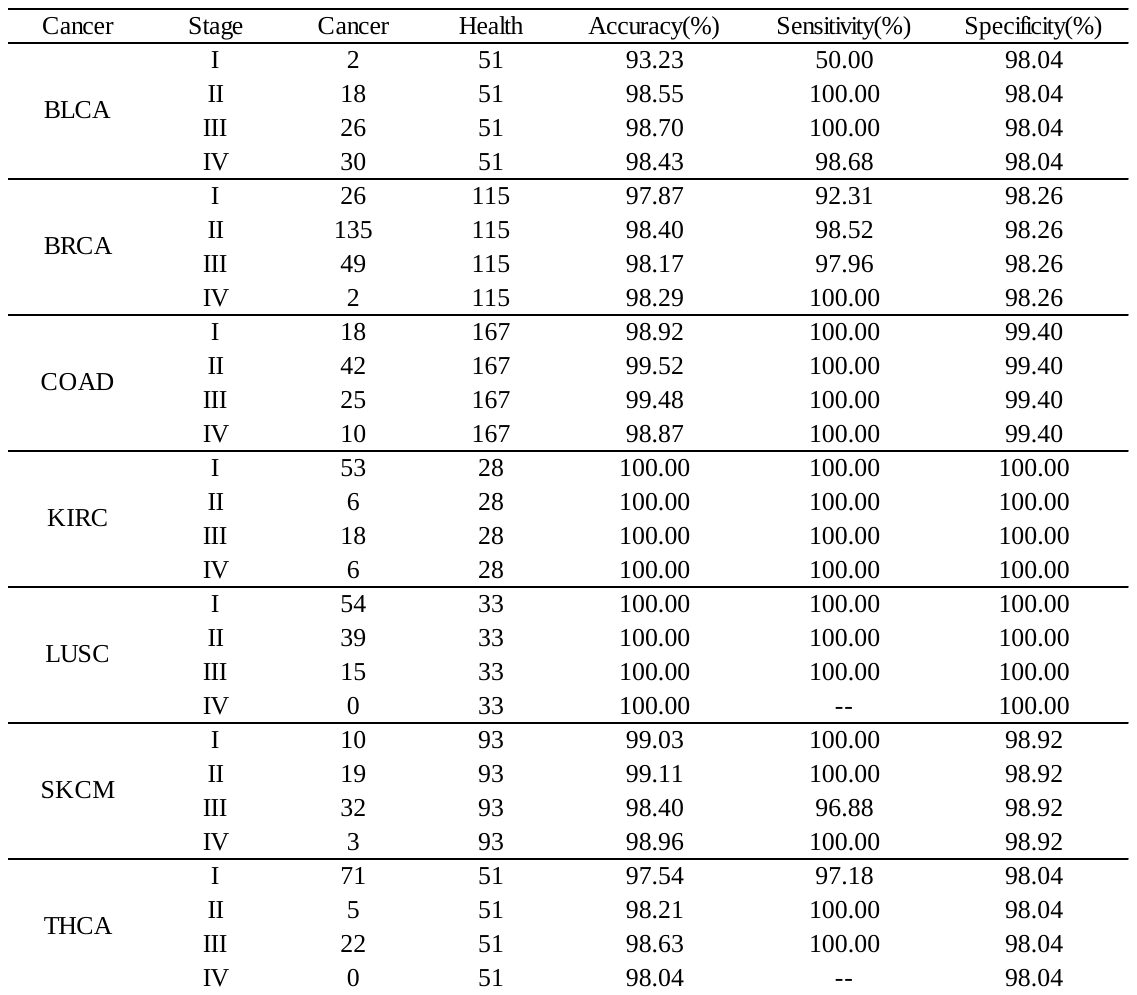

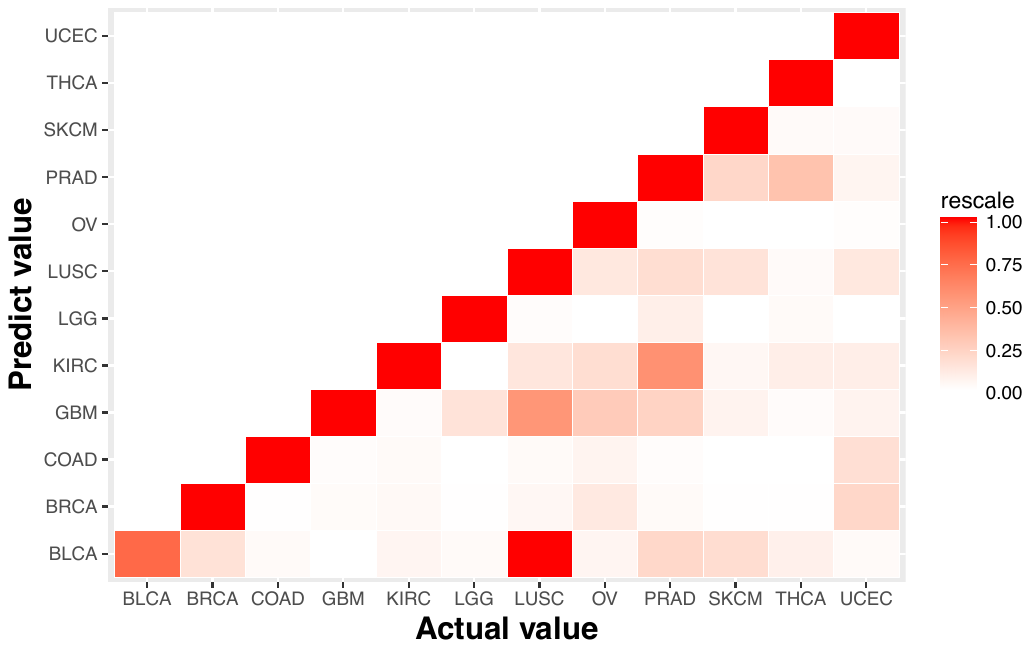

**Supplementary Figure 3 | Confusion matrix of mixture model two-way judgment.**

The misjudgment between LUSC and BLCA is most obvious, followed by LUSC and GBM, PRAD and KIRC.

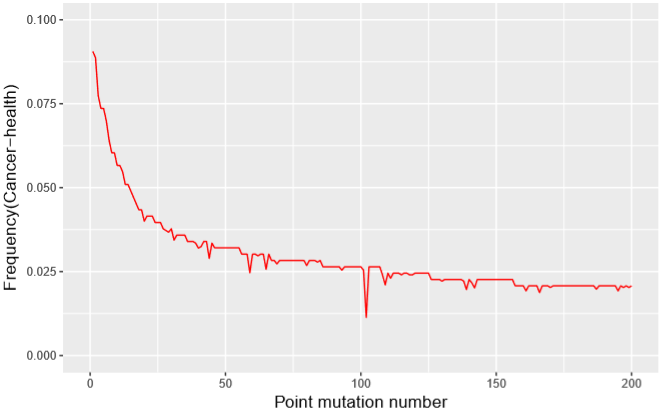

**Supplementary Figure 4 | The point mutation of uterine corpus endometrial carcinoma(UCEC).** Horizontal coordinate is the specific point mutation and Vertical coordinate standards for the difference between cancer and normal.

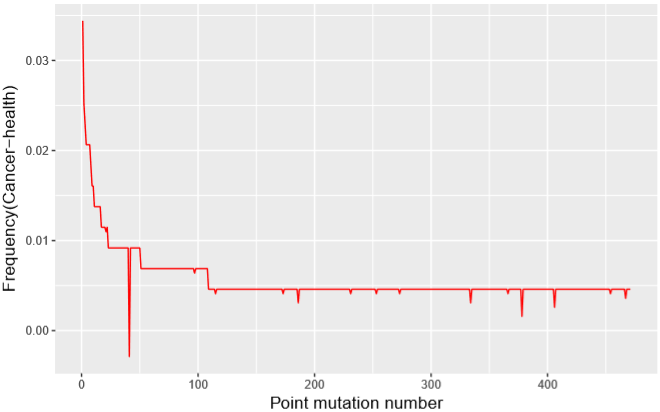

**Supplementary Figure 5 | The point mutation of ovarian carcinoma (OV).** Horizontal coordinate is the specific point mutation and Vertical coordinate standards for the difference between cancer and normal.

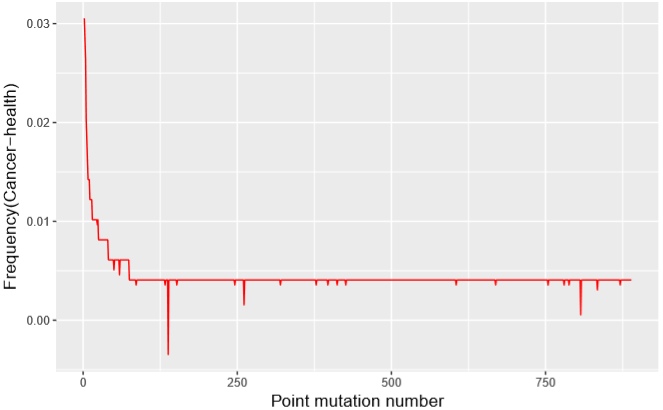

**Supplementary Figure 6 | The point mutation of lung squamous cell carcinoma (LUSC).** Horizontal coordinate is the specific point mutation and Vertical coordinate standards for the difference between cancer and normal.
